# Predicting Metabolic Dysfunction Associated Steatotic Liver Disease Risk Using Patient-Derived Induced Pluripotent Stem Cells

**DOI:** 10.1101/2025.01.13.632567

**Authors:** Yuanyuan Qin, Parth Chhetri, Elizabeth Theusch, Grace Lim, Sheila Teker, Yu-Lin Kuang, Shahrbanoo Keshavarz Aziziraftar, Mohammad Hossein Mehraban, Antonio Munoz-Howell, Varun Saxena, Dounia Le Guillou, Aras N. Mattis, Jacquelyn J. Maher, Marisa W. Medina

## Abstract

**Background and Aims:** Metabolic Dysfunction Associated Steatotic Liver Disease (MASLD) is reversible at early stages, making early identification of high-risk individuals clinically valuable. Previously, we demonstrated that patient-derived induced pluripotent stem cells (iPSCs) harboring MASLD DNA risk variants exhibit greater oleate-induced intracellular lipid accumulation than those without these variants. This study aimed to develop an iPSC-based MASLD risk predictor using functional lipid accumulation assessments.

**Methods:** We quantified oleate-induced intracellular lipid accumulation in iPSCs derived from three cohorts of diverse ancestry: 1) CIRM cohort (20 biopsy-confirmed MASH cases, 2 biopsy-confirmed MASLD cases, 17 controls), 2) POST cohort (18 MASLD cases, 17 controls), and 3) UCSF cohort (4 biopsy-confirmed MASH cases, 8 controls). Lipid accumulation levels in the CIRM cohort were used to define an iPSC-based MASLD risk score, which was used to predict case/control status in the POST and UCSF cohorts.

**Results:** In all three cohorts, lipid accumulation was higher in MASLD/MASH cases vs. controls (CIRM cases vs. controls 3.32 ± 0.25 vs. 2.70 ± 0.19 -fold change, p=0.06; POST cases vs. controls 3.63 ± 0.33 vs. 2.70 ± 0.31, p=0.05; and UCSF cases vs. controls 4.39±0.46 vs. 2.03±0.20, p=0.0002). The iPSC-based MASLD risk score achieved a sensitivity of 44% and specificity of 75% in the POST cohort and 75% and 100%, respectively, in the UCSF cohort. Differences in cohort disease severity and cardiometabolic profiles may explain performance variability.

**Conclusion:** While validation in larger cohorts is needed, these findings suggest that oleate-induced intracellular lipid accumulation in subject-derived iPSCs is predictive of MASH development. Additional cellular phenotypes and donor information should be explored to improve predictive accuracy to inform MASLD surveillance and prevention strategies.

## INTRODUCTION

Metabolic dysfunction-associated steatotic liver disease (MASLD, formerly known as NAFLD) includes a spectrum of liver phenotypes, including hepatic steatosis, which may progress to metabolic dysfunction-associated steatohepatitis (MASH, formally known as NASH). MASLD increases the risk of cardiovascular disease [1], which is the primary cause of death for these individuals[2]. Despite these risks, underdiagnosis is a significant clinical issue [3]. Generation of an individual-level predictor of MASLD onset and/or progression could be used to inform targeted surveillance, justify additional screening procedures, and promote increased attention toward preventive measures. Current models to predict high-risk MASH require the onset of disease to be informative [4-6].

MASLD is heritable (with estimates up to 50%)[7, 8]; however SNP-based heritability estimates suggest a significant fraction of the genomic contribution to MASLD remains undiscovered [9]. We reported that undifferentiated induced pluripotent stem cells (iPSCs) exhibit highly reproducible and robust accumulation of intracellular lipids in response to a fatty acid challenge, and that the magnitude of this effect is greater in cell lines that carry known MASLD genetic risk variants [10]. Here we sought to determine whether functional characterization of subject-derived undifferentiated iPSCs could be used to define an individual-level MASLD risk predictor. To our knowledge, this would represent the first attempt to use a patient-derived cell-based system to guide precision medicine in MASLD disease management.

## METHODS

### CIRM iPSC MAFLD cases and controls

The CIRM MASLD case/control cohort is comprised of men and women of diverse racial and ethnic ancestries who were recruited from the Liver Clinic at the Zuckerberg San Francisco General Hospital and Trauma Center (ZSFG). A biorepository of iPSCs from these individuals was previously described [11]. Individuals with alcohol consumption (≥ 1 drink/day for women, ≥ 2 drinks/day for men), an active HIV infection, or the presence of another liver disease were excluded. <u>CIRM MASLD cases</u> (n=22) were defined by liver biopsy. <u>CIRM Healthy Controls</u> (n=20) were defined as individuals with normal (<40U/L) ALT and AST measures.

### POST iPSC MASLD cases and controls

Through the “Pharmacogenomics of Statin Therapy (POST)” project, we recruited patients from Kaiser Permanente of Northern California (KPNC) who were at high cardiometabolic risk (i.e., individuals were users of statins, a class of cholesterol-lowering drugs). <u>POST MASLD cases</u> (n=18) were defined as individuals with one or more MASLD ICD codes, a positive diagnostic measure (ALT>40 mg/dL and AST/ALT ratio <0.8; or liver biopsy or imaging on the day of diagnosis) and lack of exclusionary medications or conditions as defined within the eMERGE NAFLD algorithm [12]. <u>POST Controls</u> (n=16) were defined as individuals without a MASLD diagnosis, normal ALT and AST (max <40 mg/dL), BMI<30, and no T2D diagnosis. The BMI and T2D exclusions were intended to mitigate the inclusion of undiagnosed MASLD cases. iPSCs were established as we previously described [13].

### UCSF iPSC MASLD cases and controls

<u>UCSF MASLD cases</u> (n=4) were previously identified as familial MASH patients, of which three are related [14]. <u>UCSF MASLD Controls</u> (n=8) were identified from the Coriell Institute as healthy donors. Fibroblasts from cases and controls were reprogrammed on feeders to iPSCs [15].

The clinical and demographic characteristics of the cohorts are shown in **Tables S1-S4**. Informed consent was obtained for the creation and distribution of all iPSCs, and studies were performed with IRB approval of ZSFG, KPNC and UCSF.

### Quantitation of intracellular lipid accumulation

iPSCs were cultured in mTeSR1 (Stem Cell Technologies) and grown at 5% O2, 5% CO_2_. Oleate-induced intracellular lipid accumulation was quantified as previously described [10]. Briefly, cells were cultured in BSA-free HCM bullet kit media (Lonza) containing 100μM oleate conjugated to BSA, or fatty acid-free BSA as a negative control for 24 hours and stained with 100ug/mL Nile Red. Fluorescence was quantified on the BD FACS LSRFortessa using the Alexa Fluor 488 and PE filters with 10,000 gated events measured. FACS data was analyzed using FloJo software.

### Generation of an iPSC-based MASLD risk score

Binary logistic regression followed by receiver operating characteristic (ROC) curve analysis was used to identify the oleate-induced lipid accumulation fold change threshold that maximizes Sensitivity-(1-Specificity) (i.e., True Positive Rate - False Positive Rate) using the CIRM iPSC dataset. The AUROC was calculated to evaluate how well iPSC lipid accumulation fold change classified the MASLD disease status of donor individuals. A similar logistic regression analysis was performed on the combined CIRM+POST+UCSF dataset (N=88).

## RESULTS

### Lipid quantification in CIRM iPSCs and creation of an iPSC-based MASLD risk score

In the 42 CIRM iPSC cell lines oleate treatment led to a 3.02 ± 0.16 (ave ± SE fold change from BSA, **Fig. 1A**) increase in intracellular lipid levels, with values increasing in iPSCs from every donor. There was a trend (p=0.06 two-tailed t-test) toward increased intracellular lipid accumulation in iPSCs from the MASLD cases (3.32 ± 0.25 fold change from BSA) vs. controls (2.70 ± 0.19 fold change from BSA), **Fig. 1B**. Using logistic regression, we found that a threshold of at least a 3.33-fold change in oleate-induced intracellular lipid accumulation best separated MASLD cases from controls.

**Fig. 1.**
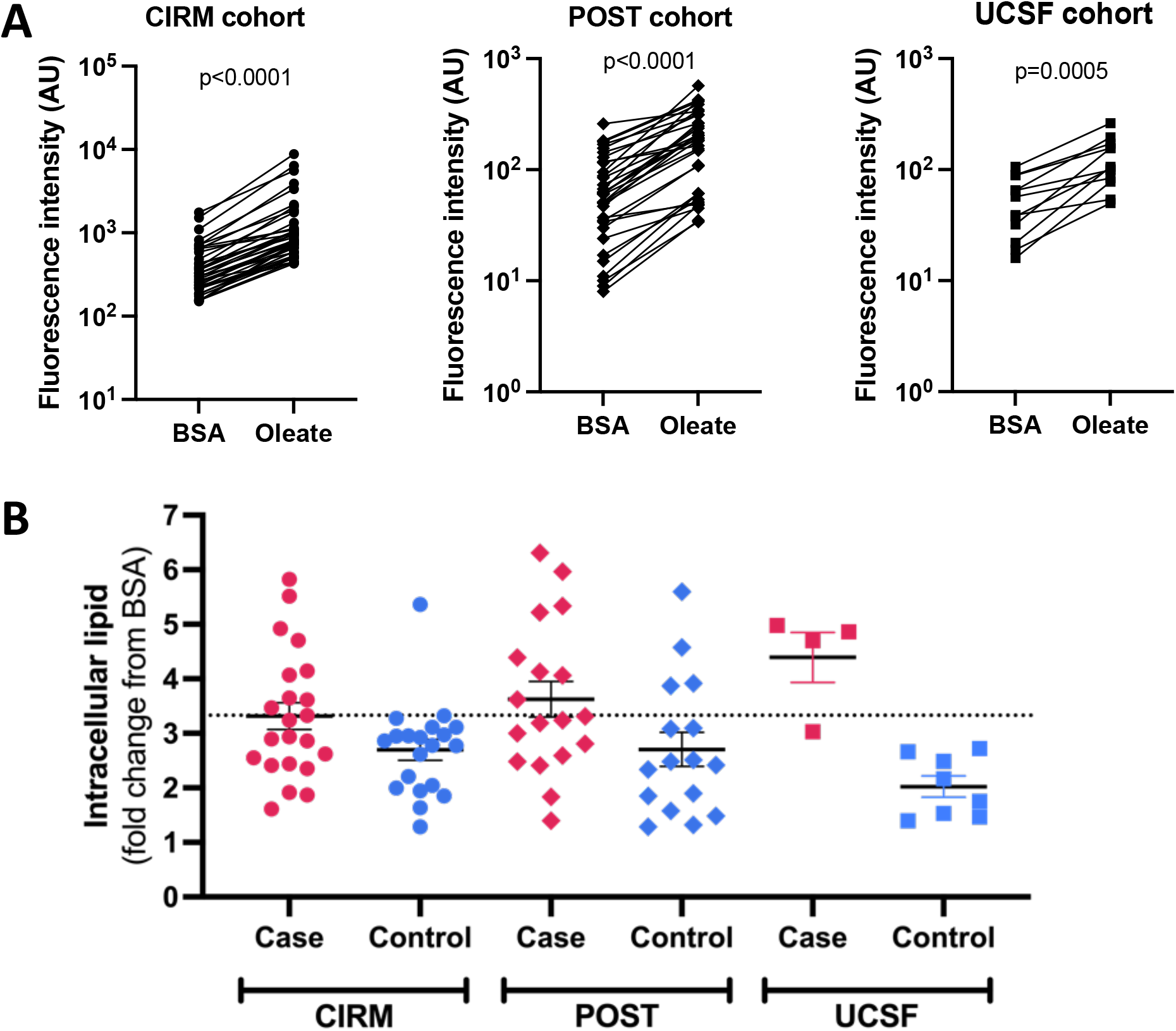
Oleate-induced intracellular lipid levels in iPSCs from three cohorts of MASLD cases and controls. iPSCs from the CIRM (n=42), POST (n=34), and UCSF (n=12) cohorts were incubated with either 100μM oleate or BSA control for 24hr after which cells were stained with Nile Red and fluorescence levels quantified by FACS. **A)** Raw fluorescence values of BSA vs. oleate-treated iPSCs. P-values were calculated using a paired Student’s t-test. **B)** Comparison of oleate-induced intracellular lipid accumulation in all three cohorts. Dashed line indicates the value of oleate-induced lipid accumulation used to define the iPSC-based MASLD risk score.

### Application of the iPSC-based MASLD risk score to the POST and UCSF cohorts

In the 35 iPSCs from the POST cohort, we observed a 3.19 ± 0.24 fold increase in intracellular lipid accumulation in response to oleate (**Fig. 1A**). There was greater intracellular lipid accumulation in the POST MASLD cases (3.63 ± 0.33) versus controls (2.70 ± 0.31), p=0.05, **Fig. 1B**. In the 12 iPSCs from the UCSF cohort, oleate-induced intracellular lipid levels rose on average 2.81 ± 0.39 fold (**Fig. 1A**), which was higher in MASH cases than controls (4.39±0.46 vs. 2.03±0.20 fold change, p=0.0002, **Fig. 1B**). Next, we tested the ability of the 3.33-fold change threshold value to correctly predict MASLD case vs. control status in the two cohorts calling cell lines with oleate-induced lipid accumulation fold change ≥3.33 as “cases” and those with values < 3.33 as “controls”. This score had a 44% sensitivity and 75% specificity to predict disease status in the POST cohort, and a 75% sensitivity and 100% specificity in the UCSF cohort.

As a secondary analysis, we recalculated the iPSC lipid accumulation fold change using all three cohorts. Application of this threshold value (3.19) to each of the cohorts individually resulted in sensitivity between 50-75%, specificity between 75-100%, precision between 73-100% and accuracy between 67-92% (**Table S5**).

### Reproducibility of the iPSC-based risk score

To evaluate the reproducibility of the iPSC-based risk score measurement we quantified the variation across technical replicates (13 iPSC lines measured in quadruplicate via FACS) and biological replicates (4 iPSC lines run on different days). Technical replicates had an average CV of 2.22 ±0.46%, while biological replicates had an average CV of 22.3±4.3% (**Fig. 2**).

**Fig. 2.**
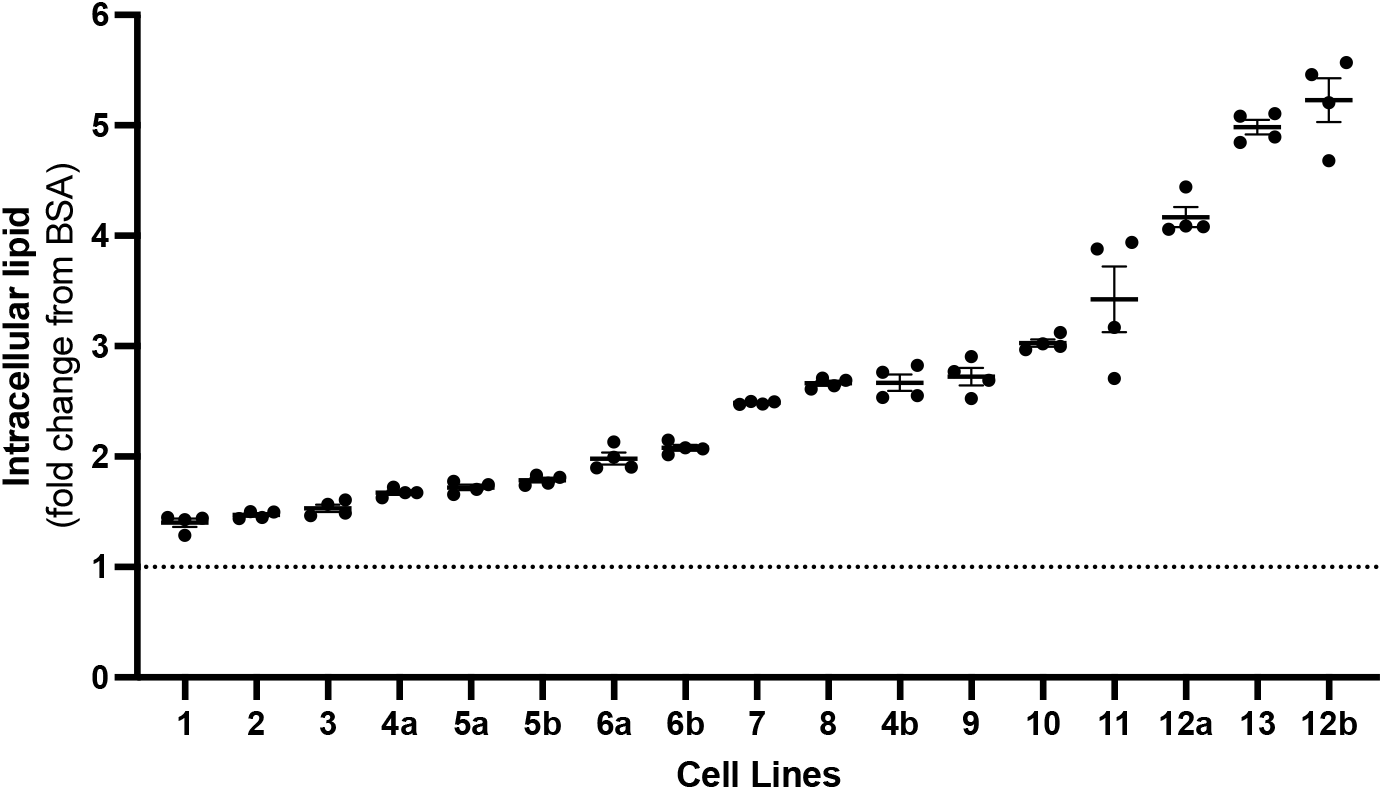
Replicate values of oleate-induced intracellular lipid accumulation. Oleate-induced intracellular lipid accumulation was measured in quadruplicate (technical replicates) by FACs in iPSCs from 13 different donors. Of these, four lines (indicated as “a” or “b”) were measured twice on different days (biological replicates).

## DISCUSSION

MASLD is a significant public health challenge, with an estimated prevalence of 30-40% among US adults [16]. The high prevalence underscores the need for innovative approaches to mitigate MASLD-related morbidity and mortality. Our study sought to address gaps in MASLD management by investigating whether the functional assessment of patient-derived iPSCs could predict individual risk of MASLD based solely on cellular data. Such a model could identify high-risk individuals prior to disease onset, providing a potential avenue for earlier intervention.

There are many existing screening algorithms to identify individuals likely to have MASLD prior to confirmation with imaging or biopsy [17-22]. However, many models use data not routinely collected in clinical settings (e.g., plasma insulin levels or waist circumference), and algorithms that do not use these data have relatively poor predictive ability [20, 21]. Even advanced machine learning algorithms and multi-omics-based approaches are largely designed to identify individuals who already have MASLD [4-6, 18], limiting their utility for proactive risk stratification. In contrast, our iPSC-based approach queries the phenotypic impact of genetic risk, offering the potential to detect disease predisposition before clinical manifestations emerge.

Advancements in technology allow for the efficient generation of patient-derived iPSCs and established protocols to differentiate them into hepatocyte-like cells (iPSC-Heps). While iPSC-Heps are valuable for studying disease mechanisms, they are costly, time-intensive, require specialized expertise, and often result in heterogeneous cultures [23]. In contrast, patient-derived iPSCs retain donor-specific genetic traits, exhibit self-renewal, and can be cryopreserved, making them a versatile and scalable cellular model. Our findings that iPSCs from MASLD/MASH cases show greater oleate-induced lipid accumulation align with previous reports, where iPSCs with MASLD genetic risk variants and iPSC-Heps from MASLD cases demonstrated higher cellular steatosis than those without these risk variants or from healthy controls [10, 11].

The CIRM iPSC-based MASLD risk score had relatively strong predictive performance in the UCSF cohort but was weaker in the POST cohort, which may be due to inherent differences in the cohorts. The CIRM and UCSF cases were either entirely or mostly MASH, while the POST cases were less severe. CIRM and UCSF cases were biopsy-defined, while POST disease status was inferred. The CIRM controls were screened to be free from liver disease, while the POST controls algorithm relied on the absence of values within the electronic health record. Lastly, all POST subjects used statins, which may reduce MASLD risk [24]. Future efforts will be needed to validate a cell-based risk assessment tool in diverse ancestry cohorts with more similar disease characteristics and evaluate whether additional information (e.g., donor characteristics, genetics, other cell-based measures, etc.) can improve the prediction performance.

In summary, while our iPSC-based model represents a promising tool for assessing MASLD genetic risk, its predictive performance varies depending on cohort characteristics. Refining these models and integrating additional layers of data could enhance their utility for early MASLD risk stratification, offering a pathway toward improved prevention and management strategies.

## Abbreviations

ALT: alanine aminotransferase
AST: aspartate aminotransferase
BMI: body mass index
BSA: bovine serum albumin
CV: coefficient of variation
FACS: fluorescence-activated cell sorting
ICD: international classification of diseases
iPSC: induced pluripotent stem cell
iPSC-Hep: induced pluripotent stem cell-derived hepatocyte-like cell
MASLD: metabolic dysfunction associated steatotic liver disease
MASH: metabolic dysfunction steatohepatitis
SNP: single nucleotide polymorphism
T2D: Type 2 Diabetes

## Acknowledgements

We would like to thank the following individuals for their assistance in cell culturing, sample preparation, and organization: Julia Su, Leela Venkatesan, Sai Chelluri, and Yuqing Zhang. We thank all study participants without whom this work would not be possible. Samples were analyzed using the UCSF MLK Core Research Facility.

## REFERENCES

1. Bhatia, L.S., et al., Non-alcoholic fatty liver disease: a new and important cardiovascular risk factor? Eur Heart J, 2012. 33(10): p. 1190–200.

2. Chalasani, N., et al., The diagnosis and management of nonalcoholic fatty liver disease: Practice guidance from the American Association for the Study of Liver Diseases. Hepatology, 2018. 67(1): p. 328–357.

3. Petta, S., et al., Prevalence and severity of nonalcoholic fatty liver disease by transient elastography: Genetic and metabolic risk factors in a general population. Liver Int, 2018. 38(11): p. 2060–2068.

4. Njei, B., et al., An explainable machine learning model for prediction of high-risk nonalcoholic steatohepatitis. Sci Rep, 2024. 14(1): p. 8589.

5. Docherty, M., et al., Development of a novel machine learning model to predict presence of nonalcoholic steatohepatitis. J Am Med Inform Assoc, 2021. 28(6): p. 1235–1241.

6. Ghandian, S., et al., Machine learning to predict progression of non-alcoholic fatty liver to non-alcoholic steatohepatitis or fibrosis. JGH Open, 2022. 6(3): p. 196–204.

7. Loomba, R., et al., Heritability of Hepatic Fibrosis and Steatosis Based on a Prospective Twin Study. Gastroenterology, 2015. 149(7): p. 1784–93.

8. Schwimmer, J.B., et al., Heritability of nonalcoholic fatty liver disease. Gastroenterology, 2009. 136(5): p. 1585–92.

9. Vujkovic, M., et al., A multiancestry genome-wide association study of unexplained chronic ALT elevation as a proxy for nonalcoholic fatty liver disease with histological and radiological validation. Nat Genet, 2022. 54(6): p. 761–771.

10. Munoz, A., et al., Undifferentiated Induced Pluripotent Stem Cells as a Genetic Model for Nonalcoholic Fatty Liver Disease. Cell Mol Gastroenterol Hepatol, 2022. 14(5): p. 1174–1176 e6.

11. Duwaerts, C.C., et al., iPSC-derived hepatocytes from patients with nonalcoholic fatty liver disease display a disease-specific gene expression profile. Gastroenterology, 2021.

12. Namjou, B., et al., GWAS and enrichment analyses of non-alcoholic fatty liver disease identify new trait-associated genes and pathways across eMERGE Network. BMC Med, 2019. 17(1): p. 135.

13. Kuang, Y.L., et al., Evaluation of commonly used ectoderm markers in iPSC trilineage differentiation. Stem Cell Res, 2019. 37: p. 101434.

14. Mattis, A.N., et al., Modeling Familial NASH by iPSC-Hepatocytes. submission in process.

15. Yu, J., et al., Human induced pluripotent stem cells free of vector and transgene sequences. Science, 2009. 324(5928): p. 797–801.

16. Spengler, E.K. and R. Loomba, Recommendations for Diagnosis, Referral for Liver Biopsy, and Treatment of Nonalcoholic Fatty Liver Disease and Nonalcoholic Steatohepatitis. Mayo Clin Proc, 2015. 90(9): p. 1233–46.

17. Kuhn, T., et al., Anthropometric and blood parameters for the prediction of NAFLD among overweight and obese adults. BMC Gastroenterol, 2018. 18(1): p. 113.

18. Atabaki-Pasdar, N., et al., Predicting and elucidating the etiology of fatty liver disease: A machine learning modeling and validation study in the IMI DIRECT cohorts. PLoS Med, 2020. 17(6): p. e1003149.

19. Kotronen, A., et al., Prediction of non-alcoholic fatty liver disease and liver fat using metabolic and genetic factors. Gastroenterology, 2009. 137(3): p. 865–72.

20. Long, M.T., et al., Development and Validation of the Framingham Steatosis Index to Identify Persons With Hepatic Steatosis. Clin Gastroenterol Hepatol, 2016. 14(8): p. 1172-1180.e2.

21. Lee, J.H., et al., Hepatic steatosis index: a simple screening tool reflecting nonalcoholic fatty liver disease. Dig Liver Dis, 2010. 42(7): p. 503–8.

22. Bedogni, G., et al., The Fatty Liver Index: a simple and accurate predictor of hepatic steatosis in the general population. BMC Gastroenterol, 2006. 6: p. 33.

23. Godoy, P., et al., Recent advances in 2D and 3D in vitro systems using primary hepatocytes, alternative hepatocyte sources and non-parenchymal liver cells and their use in investigating mechanisms of hepatotoxicity, cell signaling and ADME. Arch Toxicol, 2013. 87(8): p. 1315–530.

24. Ayada, I., et al., Dissecting the multifaceted impact of statin use on fatty liver disease: a multidimensional study. EBioMedicine, 2023. 87: p. 104392.

